# IGF1 modulates lesional skin inflammation in checkpoint inhibitor-induced lichen planus

**DOI:** 10.64898/2026.05.27.726087

**Authors:** Noah I Hornick, Avery Billo, Rosalyn M Fey, Reed M Hawkins, Fiorinda F Muhaj, Kristen N Richards, Anisha B Patel, Jason M Schenkel, Kristen E Pauken, Amy E Moran

**Author notes:** These authors contributed equally to this work (Noah I Hornick, Avery Billo).

## Abstract

Immune checkpoint inhibitor-induced lichen planus (ICI-LP) is a cutaneous immune related adverse event (irAE) that shares key clinicopathologic features with spontaneous lichen planus (LP) but differs histologically and in the sex distribution of its incidence, and may therefore reflect a distinct tissue inflammatory state. To define the cellular programs that distinguish ICI-LP from LP, we profiled lesional skin by single cell and spatial transcriptomic approaches. We found few differences in the T cell and keratinocyte compartments between ICI-LP and LP, which shared similar inflammatory signatures. Rather, the dominant transcriptional features differentiating these two eruptions occurred within the fibroblast and myeloid cell compartments. Fibroblasts in ICI-LP were enriched for IGF1, FGF7, and androgen-response-associated programs, whereas myeloid cells exhibited amplified JAK-STAT and interferon-responsive states spanning both type I and type II interferon signatures. The potential role of androgen response in shaping lichenoid inflammation was supported by a striking loss of androgen receptor expression in lesional keratinocytes by immunohistochemistry. Furthermore, using spatial RNA and transcriptomic approaches, we identified anatomically segregated *IFNG, IL17A*, and *IL13* niches within lesional skin, suggesting that regional immune compartmentalization with differences in local immunoregulation may explain the mixed inflammatory features reported in both ICI-LP and LP. Collectively, these data indicate that ICI-LP is not simply a more inflamed form of LP, but a distinct form of the disease with more prominent inflammatory perturbations within stromal and innate immune cell populations.

## Introduction

The potential reach of immune checkpoint inhibitors (ICIs) has been limited by the frequency and severity of immune-related adverse events (irAEs)^1^. These reactions affect up to 90% of treated patients^2^, with a higher incidence in women than in men^3–5^. While irAEs can affect any organ, they are most frequent in barrier tissues, particularly the skin^6–10^. Cutaneous irAEs (cirAEs) occur across a spectrum of clinical presentations, recapitulating a broad swath of inflammatory dermatology^11–14^, and, as with irAEs generally, disproportionately affecting women^15^. Cutaneous irAEs lead to treatment interruption or discontinuation in as many as 24.3% of cases^13^. The rapid and ongoing expansion of indications and contexts for ICIs underscores the importance of better managing the growing burden of cirAEs. This need is particularly acute in dermatology, where cutaneous toxicities are common, can be diagnostically challenging, and may require modification or cessation of otherwise effective anti-cancer therapy.

ICI-induced lichen planus (ICI-LP) is a representative example of a cirAE that both resembles and distinguishes itself from an established inflammatory skin disease. It is among the more common cirAE, affecting 17% of ICI-treated patients^16^, with both clinical and histopathologic features that differentiate it from its namesake, lichen planus (LP)^17–20^. Although both eruptions share a band-like lymphohistiocytic infiltrate, acanthosis, hyperkeratosis, and dyskeratosis, ICI-LP has distinguishing histopathologic features including a CD4^+^-predominant T cell infiltrate, spongiosis, parakeratosis, and eosinophils^18,21^. More recent reports also note increased infiltration of lymphocytes into the epidermis along with increased histiocytes in the inflammatory infiltrate^22,23^. From an epidemiologic perspective, ICI-LP exhibits a reversal of the sex distribution observed in LP: while LP is more frequent in women^24–27^, ICI-LP-affected patients are equally likely to be male or female^28^, and in some cohorts are more likely to be male^13,18,29,30^. This finding goes against the more broadly-described sex differences in response to ICIs; male sex is associated with better anti-tumor response^31,32^, while female sex is associated with higher rates and severity of irAEs^3–5^. Collectively, these features suggest that distinct immunoregulatory features are likely to exist between LP and ICI-LP, and likely relate to their sex distribution. Unlike several other common cirAEs, ICI-LP also lacks a well-defined mechanistic framework that could guide targeted, steroid-sparing therapy. Because ICI-LP and spontaneous LP share a common lichenoid reaction pattern while differing in clinical context, histopathology, and sex distribution, direct comparison offers an opportunity to identify tissue programs specific to checkpoint blockade-associated disease.

Sex hormones exert a variety of effects on the adaptive immune system, culminating in a set of clinical differences in immune responses between men and women. For example, women experience more autoimmune disease^33^, while men experience higher rates of cancer^34^. Mechanistically, these impacts are often mediated by the direct actions of sex hormones themselves. Androgen receptor (AR) signaling in particular can suppress Th1 differentiation^35,36^, directly reduce IFN-γ *product*ion in activated T cells^37^, and limit antigen-specific T cell responses^38^, while 17β-estradiol signaling can promote Th1 or Th2 differentiation^39^. Recent work has demonstrated that the skin is among the tissues with the highest levels of sex-based differential gene expression^40^; androgens contribute to this by suppressing innate lymphoid cell 2 (ILC2) populations, and by altering dendritic cell activation and downstream T helper polarization^41,42^. The reversal in sex distribution observed in ICI-LP therefore raises the possibility that local hormone-responsive programs may shape disease phenotype. More broadly, however, the mechanisms that distinguish ICI-LP from spontaneous LP remain poorly defined at the level of the tissue microenvironment.

While most existing literature supports a dominant role for IFN-γ*-driven* Th1 inflammation in LP^43,44^, there is significant evidence for additional complexity, with putative roles for multiple other T helper subtypes^45–47^. From the clinical perspective, it is notable that targeting of the Th1 response in LP has been more successful than targeting other T cell response types. IFN-γ *signali*ng through JAK2/STAT1 is central to keratinocyte damage in LP^43^, and the efficacy of JAK inhibitors in LP has paralleled their ability to impact JAK2, with decreased rates of response seen when targeting is directed toward pathways with greater roles in Th2 or Th17 inflammation^48–51^. At the same time, inflammatory skin disease is increasingly understood to be shaped not only by lymphocytes, but also by stromal and innate immune populations that help organize local tissue immune states. While lichenoid inflammation is often framed primarily through T cell-mediated keratinocyte injury, whether fibroblast and myeloid cell populations contribute to the divergence between LP and ICI-LP remains unknown.

Although data regarding ICI-LP is more limited, several notable recent findings support the emergence of Th2 features through a dominant IFN-γ*/*Th1 milieu that are potentially more substantial than those seen in LP. A recent cytokine analysis that included serum, tape-stripped skin, and skin biopsy samples found that in ICI-LP, IFN-γ, IL-13, and IL-17A are all significantly increased versus normal skin^52^. A study comparing ICI-induced eczematous eruptions, ICI-LP, and spontaneous lichenoid skin disease found that Th2-polarized T cells were overrepresented in ICI-LP compared to spontaneous lichenoid disease, and on par with levels seen in eczematous eruptions^28^. Lastly, and perhaps most significantly, ICI-LP has been reported to respond to IL-4/IL-13 inhibition with dupilumab^53^. These findings suggest that ICI-LP may arise through tissue immune programs that overlap with, but are not identical to, those operating in spontaneous LP. The apparent coexistence of Th1-, Th2-, and Th17-associated features also raises the possibility that regional immune compartmentalization within lesional skin contributes to the heterogeneous inflammatory phenotypes attributed to both LP and ICI-LP.

We therefore performed a comparative transcriptional and spatial analysis of LP and ICI-LP lesional skin to define the cellular compartments and inflammatory programs that distinguish these related conditions. Given the unusual sex distribution of ICI-LP and the known immunoregulatory effects of sex hormones in skin, we also asked whether hormone-responsive programs might contribute to these differences.

## Results

### Androgen receptor expression is diminished in inflamed epidermis and dermal stroma in LP and ICI-LP

Sex differences in cancer pathogenesis, progression, treatment response, and toxicity are complex and multifaceted, comprising numerous influences ranging from genetic to environmental^54-56^. Given the unusual sex distribution of ICI-LP relative to LP and the emerging literature on hormone-responsive immune programs, we first examined androgen receptor (AR) expression in lesional skin by immunohistochemistry^35-42,57^. While AR was readily detectable in cutaneous keratinocytes and fibroblasts in normal controls (**Fig 1A, top**), expression was strikingly absent from keratinocytes in more inflamed regions of epidermis in LP (**Fig 1A, center**) and ICI-LP (**Fig 1A, bottom**). Fibroblast expression of AR, however, was still visible in the dermis of both ICI-LP and LP samples. These data demonstrate a loss of AR expression in both LP and ICI-LP stroma (**Fig 1B**), focused in the areas immediately surrounding the inflammatory infiltrate.

**Figure 1:**
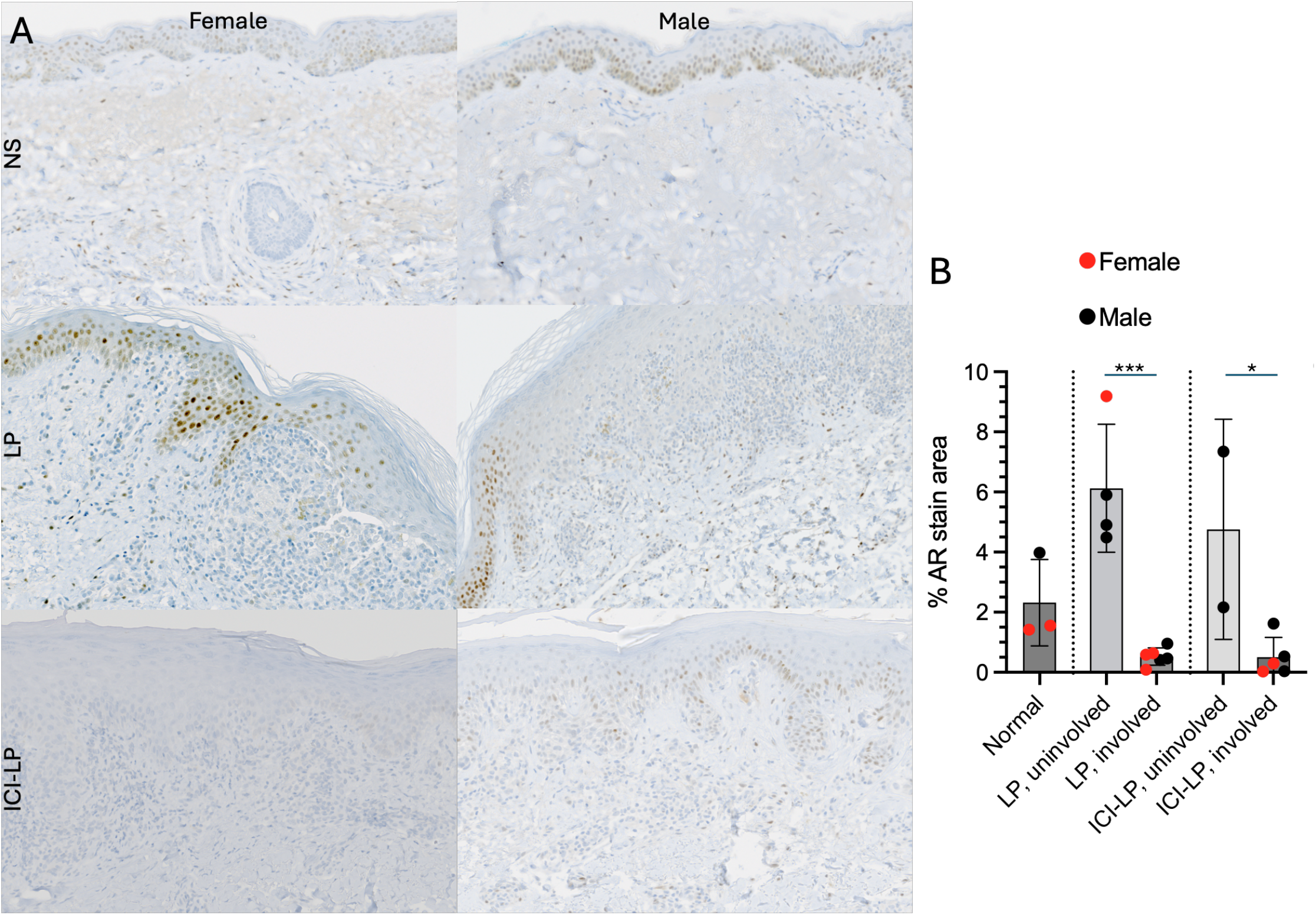
AR is decreased in lesional epidermis in LP and ICI-LP. (A)IHC for AR in normal skin (NS,top), lesional skin from LP (center),and lesional skin from ICI-LP(bottom) in representative Female (left)and Male (right)patient.(B)Quantification of staining in samples represented by(A).*p<0.05;***,p<0.001

### Single cell profiling identifies similar lesional constituents but selective expansion of myeloid cells and proliferating T cells in ICI-LP

To determine which tissue compartments underlie these histologic differences and the altered pattern of AR expression, we next performed single-cell RNA sequencing of lesional samples from patients with LP and ICI-LP. For these investigations, we restricted our analysis to male patients to reduce variability in factors impacting baseline androgen levels (for patient characteristics see **Table S1**). The specimens consisted of residual tissue from clinical biopsy specimens and had been formalin fixed and paraffin embedded. To ensure that this processing did not impact our results, we compared sequencing results from these samples with those obtained during prior studies using fresh tissue^58^ and found no processing-specific differences in cell types identified, nor any UMAP clusters specific to one processing method among major skin cell types (**Fig S1**).Initial unsupervised clustering and comparison of cell populations between ICI-LP and LP patient skin demonstrated largely similar cell types (**Fig 2A, Fig S2A-B**), with significantly greater proportions of proliferating T cells and myeloid populations in ICI-LP (**Fig 2B**). Using pseudobulk analysis, differential expression (DE) analysis at a whole-sample level revealed elevated *IGF1, FGF7*, and to a lesser extent *SRPX* in ICI-LP samples as compared to LP (**Fig 2C**). We examined cluster-by-cluster expression of these genes to determine the cell types driving this gene expression and found significant overexpression of all three mRNA species to be concentrated in fibroblasts (**Fig 2D, Fig S2C**). We next assessed signaling pathway activation using PROGENy pathway signatures^59^, and found that JAK-STAT signaling is the most differentially activated pathway on a whole-tissue level between ICI-LP and LP (**Fig 2E**), supporting a dominant role for Th1 immunity in ICI-LP. While previous studies have also demonstrated marked Th1 activity in ICI-LP, this was accompanied by elevations in cytokines and cell populations associated with Th2 and Th17 activation^23,28,60^. Moreover, although ICI-LP and LP shared a broadly similar cellular architecture, the dominant transcriptional differences between them localized to fibroblast and myeloid compartments rather than to T cells or keratinocytes (**Fig 2F**). Specifically, while elevated JAK-STAT activation was evident in ICI-LP myeloid cells (and to a lesser extent T cells), a different response was found in fibroblasts (**Fig 2F**). Specifically evaluating the pathways to which IGF1, FGF7, and SRPX contribute revealed that the androgen response pathway, to which IGF1 is a contributor^59^, is markedly increased in ICI-LP fibroblasts (**Fig 2F**). These findings highlight key similarities and differences between ICI-LP and LP. While ICI-LP has a greater enrichment in proliferating T cells, more substantial changes were observed in fibroblasts and myeloid cells than the T cell compartment. ICI-LP seemed more proinflammatory than LP, particularly in the myeloid compartment, while fibroblasts showed upregulated *IGF1, FGF7*, and *SRPX*. These findings are consistent with the notion that ICI-LP has both clinical and histopathologic features that distinguish it from LP, and these differences also manifest in the transcriptional landscape of stromal compartments within these lesions.

**Figure 2:**
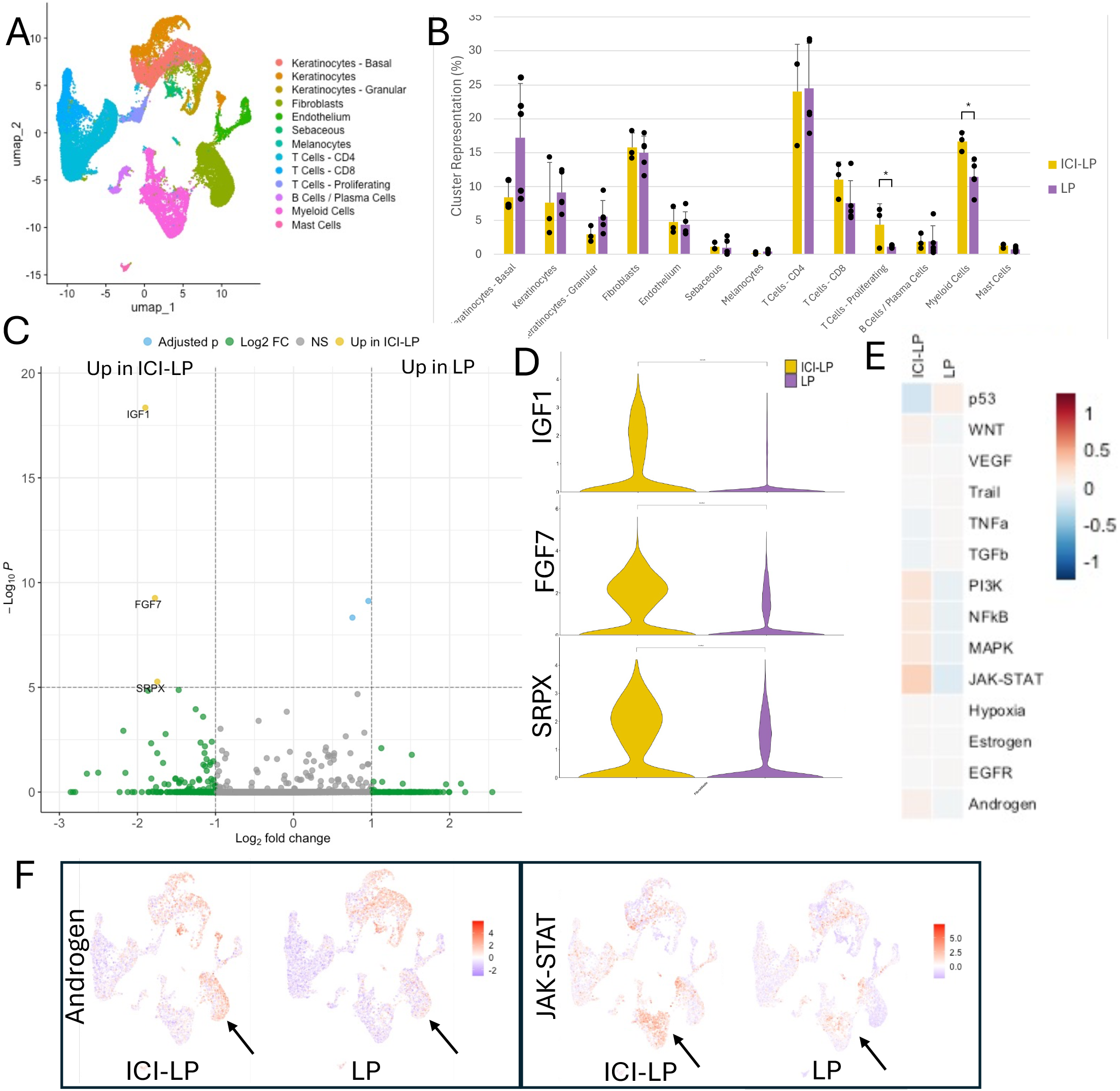
ICI-LP lesions are similar to LP in cell type representation; myeloid cells are overrepresented in ICI-LP. (A) UMAP plot of scRNAseq of lesional skin of LP and ICI-LP. (B) Contribution of UMAP clusters to total composition of cells isolated on a per-sample basis comparing ICI-LP to LP. (C) DiDerentially expressed genes between ICI-LP and LP lesions based on pseudobulking.Threshold for annotation is Log_2_ fold change > 1 or < -1 and -log_10_ p > 5. (D) Expression of IGF1, FGF7, and SRPX by Fibroblast UMAP cluster. (E) Comparison of pathway activation in ICI-LP and LP samples using PROGENy. (F) Androgen (left) and JAK-STAT (right) pathway activation in UMAP clusters. Arrows denote fibroblasts (left) and myeloid cells (right). *, p<0.05; **, p<0.01; ***, p<0.001; ****, p<0.0001.

### Fibroblasts are a major distinguishing compartment in ICI-LP and exhibit enriched IGF1/FGF7 and androgen response programs

Because IGF1, FGF7, and SRPX expression localized primarily to fibroblasts, we next pursued a focused analysis of fibroblast states in ICI-LP and LP. Previous work demonstrated that fibroblasts can support Th2-directed immune responses in the skin^61^. Our initial subclustering of lesional fibroblasts revealed populations with distinct transcriptional identities; these were either dominated by gene programs associated with tissue location (papillary or reticular) or by phenotype^61,62^ (**Fig 3A**). The identified clusters made up similar proportions of fibroblasts in ICI-LP and LP skin, although Superficial/Papillary Dermal FRC-Like fibroblasts were decreased in ICI-LP (**Fig 3B**). While some populations were clearly defined by location or by phenotype alone, there were clusters that overlapped in role-defining gene expression (**Fig 3C**). DE analysis confirmed elevated expression of *IGF1* in ICI-LP fibroblasts with marked elevation of *ADAMDEC1* in fibroblasts derived from LP lesions (**Fig 3D**). Expression of *ADAMDEC1* and *CXCL9* have previously been identified as activation markers associated with “disease-adapted” FRC-like fibroblasts, including in lichenoid drug reactions^62^. Interestingly, while expression of these genes is indeed primarily evident in FRC-like fibroblast populations, *CXCL9* is expressed in both ICI-LP and LP, while *ADAMDEC1* is relatively restricted to fibroblasts from LP (**Fig 3E**). Next, to examine modules of coordinated gene expression that might further differentiate transcriptional states among fibroblasts in ICI-LP and LP, we performed topic modeling using latent Dirichlet allocation (LDA)^63^. This process defines randomly-numbered topics – sets of genes whose tendency to be co-expressed within cells can be leveraged to identify transcriptional patterns within single-cell data. The topic most overexpressed in ICI-LP-associated fibroblasts (**Fig 3F**) included *IGF1, FGF7, TNC* (reported to enhance cutaneous inflammation in murine imiquimod-mediated dermatitis^64^), and other genes associated with IGF1 signaling (*IGFBP2* and *IGFBP4*^65^) and TGF β signaling (*NR4A1*^66^, *THBS1*^67^, *CEMIP*^68^, *GREM1*^69^). Pathway activation in lesional fibroblasts demonstrated increased androgen signaling in ICI-LP, which appeared to affect all subpopulations of fibroblasts (**Fig 3G**,**H**). These data are consistent with an elevated androgen response in ICI-LP, a key difference between ICI-LP and LP.

**Figure 3:**
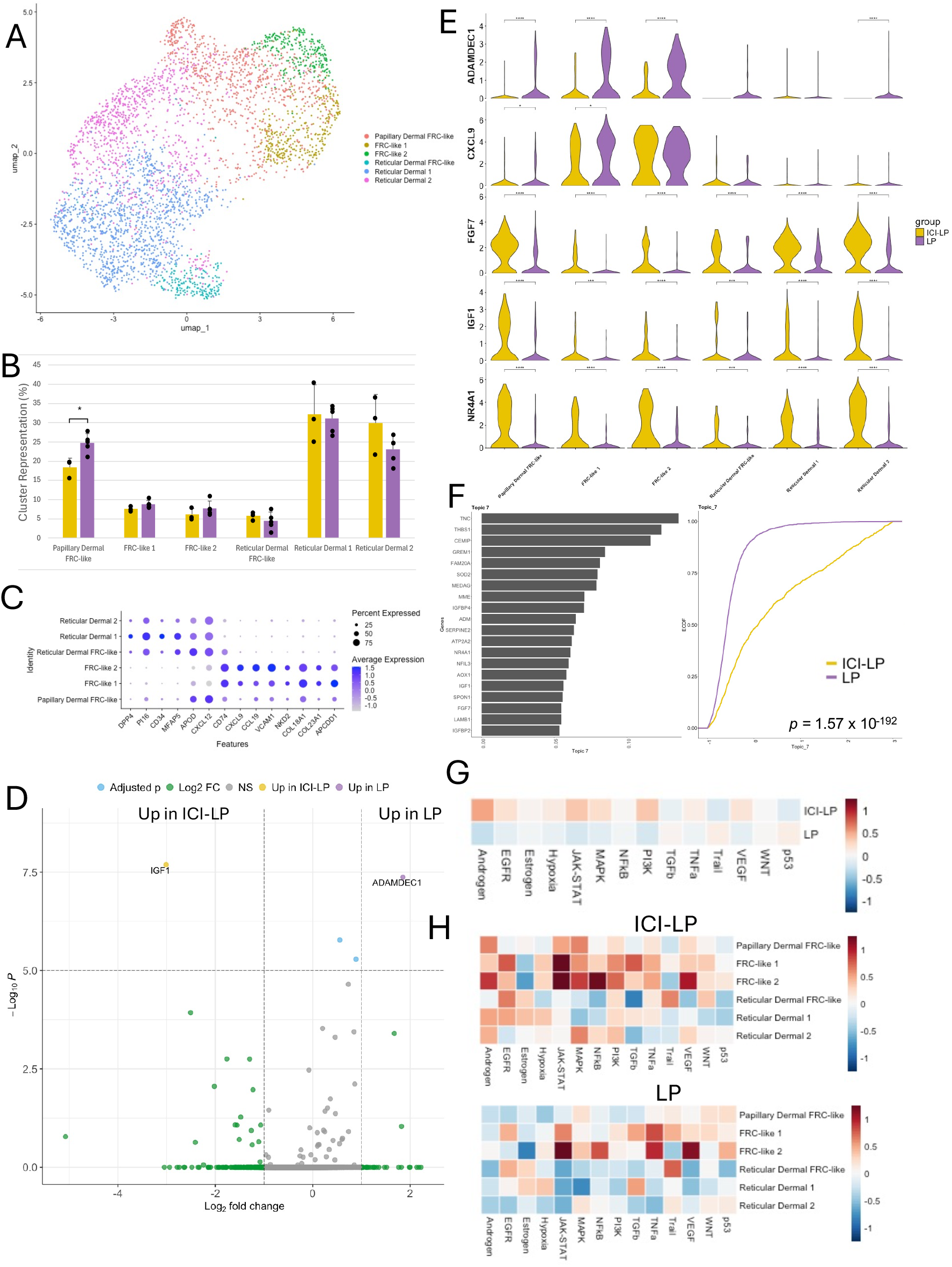
ICI-LP fibroblasts exhibit increased androgen response focused on IGF1. (A) UMAP plot of fibroblast subclustering. (B) Contribution of UMAP clusters to total composition of fibroblasts, comparing ICI-LP to LP. (C) Dot plot of marker genes used in cell type annotation in B. (D) DiDerentially expressed genes between ICI-LP and LP lesions based on pseudobulking. Threshold for annotation is Log_2_ fold change > 1 or < -1 and -log_10_ p > 5. (E) Expression of ADAMDEC1, IGF1, and NR4A1 by UMAP cluster. (F) Topic 7: diDerentially expressed gene set between ICI-LP and LP based on LDA analysis. Left: bar plot for top-ranked genes; Right: empirical cumulative distribution function (ECDF) of cells per topic by condition (ICI-LP vs LP). NB: the lower line indicates increased topic gene expression in ECDF plots. (G) Comparison of pathway activation in ICI-LP and LP fibroblasts using PROGENy. (H) PROGENy pathway activation by subcluster. *, p<0.05; **, p<0.01; ***, p<0.001; ****, p<0.0001.

### Keratinocytes are similar in ICI-LP and LP

Keratinocytes have been shown to actively contribute to the inflammatory response in multiple spontaneous inflammatory diseases of the skin^70,71^ and our analysis revealed striking loss of AR expression by immunohistochemistry (**Fig 1A**). Therefore, we asked whether keratinocyte states differed between ICI-LP and LP. Comparison of keratinocytes overall and by subclustering revealed remarkably little variability between the two conditions. Subpopulation representation and DE analysis revealed only variability in representation of follicles (increased PHLDA1, a follicular marker, in ICI-LP) and adnexa (increased representation of eccrine keratinocytes in ICI-LP) (**Fig S3B-D**), which seemed more likely to be related to biopsy site and variable sampling than a biological difference.

### Myeloid populations show elevated IFN response signatures in ICI-LP compared to LP

Androgen upregulates *IGF1* expression and signaling^72^, and both androgen and IGF1 signaling can promote JAK/STAT activity^73,74^ – the pathway elevation seen primarily in myeloid cells in the initial whole-tissue analysis (**Fig 2F**). We therefore next turned our attention to myeloid cell populations to evaluate whether androgen signaling or fibroblast-produced IGF1 and FGF7 might contribute to this elevated JAK/STAT activity or to their overrepresentation as a proportion of the ICI-LP inflammatory infiltrate (**Fig 2B**). Subclustering of myeloid cells revealed expected cutaneous populations. These included Langerhans cells and types 1 and 2 conventional dendritic cells (cDC1 and cDC2, respectively) as well as mature dendric cells enriched in immunoregulatory molecules (mregDCs) distinguished by *LAMP3* and *CCR7* expression. Also present were macrophages expressing either *CD169* or *C1q*; and a group of cells expressing *CCR2, ITGAX, S100A8, S100A9*, and *VCAN* that were transcriptionally similar to monocytic myeloid-derived suppressor cells (mMDSC^75^) (**Fig 4A**). ICI-LP and LP contain similar proportions of each population, with CD169^+^ macrophages being overrepresented and Langerhans cells underrepresented in ICI-LP (**Fig 4B**). An examination of differential gene expression in these cells yielded a cluster of genes upregulated in ICI-LP samples including *IFITM1, ISG15, ISG20, PLAC8, S100A9*, and *S100A12* (**Fig 4C**). Genes identified as elevated in LP (**Fig 4C**) seemed likely related to the difference in Langerhans cell representation (**Fig 4B, Fig S4A**). The enrichment of DEGs in ICI-LP was consistent with our prior observation (**Fig 2F**) of increased activation of JAK-STAT in myeloid cells given the inclusion of *IFITM1, ISG15*, and *ISG20*, all of which contribute to the PROGENy JAK-STAT score^59^. Indeed, employing pathway analysis on these populations demonstrated increased JAK-STAT activity across all myeloid populations in ICI-LP compared to LP (**Fig 4D,E**). *IFITM1, ISG15*, and *ISG20* mirrored this finding and were expressed across all subpopulations with the exception of Langerhans cells. *S100A9* and *S100A12* expression, however, was largely restricted to the mMDSC cluster (**Fig S4B**). Given the similar fold changes and p values of the genes overexpressed in ICI-LP, we again turned to topic modeling to examine whether these were members of a larger, potentially co-regulated, gene expression program that could be empirically determined. *S100A9* and *ISG15* were both identified as major contributors to the identity of Topic 2 (**Fig 4F**), which also contained *S100A8* and *MX2*, was primarily expressed in the mMDSC cluster (**Fig S4C**), and was overexpressed in ICI-LP. As both *ISG15* and *MX2* are associated with interferon response, we were intrigued to note that the topic most overexpressed in ICI-LP, Topic 12, was also composed of genes typical of interferon response, including *MX1, IFITM3, OAS2*, and *OAS3* (**Fig 4F**). It also included *VCAN* and was primarily expressed in the mMDSC cluster (**Fig S4C**). Because *MX1* is more characteristically induced by Type I than Type II interferon^76^, we assessed the activity of IFNα (Type I interferon) and IFNγ (Type II interferon) signaling using established gene expression signatures of interferon response^77,78^. We found increased expression of both Type I and Type II response in ICI-LP-derived over LP-derived myeloid cells, with a slightly greater increase in Type I response (**Fig 4G**). Collectively, this evidence supports the increase in JAK/STAT signaling (downstream of interferons) observed in ICI-LP and underscores a dominant effect of interferon signaling in lesional ICI-LP and LP skin.

**Figure 4:**
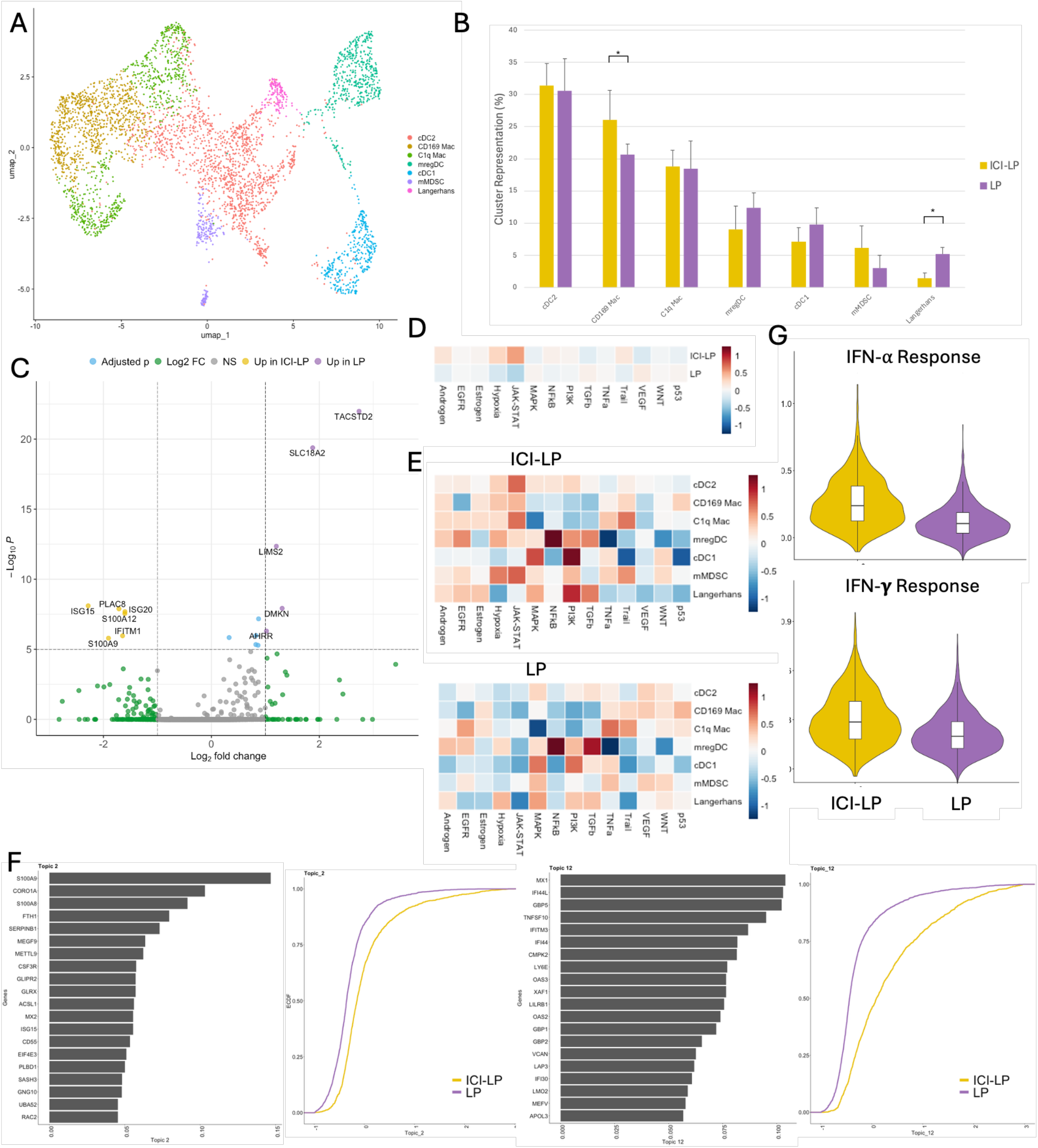
ICI-Lp myloid populations show elevate type 1 interferon response. (A)UMAP plot of myeliod cell subclustering.(B) Contribution of UMAP clusters tot total composition of myeliod cells,comparing ICI-LP to LP.(C) Differentially expressed ngenses betweenICI-LP and LP myeliod cells,pseudobulking.Threshold for annotation is Log_2_ fold change >1or<-1and -log_10_p>5.(D) Comparision of pathyway activation in ICI-LPmyeliod cells using PROGENy.(E) PROGENy pathyway activation by subcluster.(F)Differentially expressed topics expressed topics between ICI-LPand LP based on LDA analysis.Left ofeach pair: bar plot for top-ranked genes;Right:empirical cumlative distribution function (ECDF)of cells per topic by condition (ICI-LPvs LP).(G) Modules scores of IFN-αand IFN-γ in my eliod cells.*,p<0.05.

### T cells are broadly similar between ICI-LP and LP, with limited transcriptional differences

Although fibroblast and myeloid populations emerged as the dominant compartments distinguishing ICI-LP from LP, lichenoid inflammation is traditionally thought of as a T cell-mediated process. Therefore, we next asked whether T cell states also differed between the two conditions. Subclustering and annotation of T cells yielded a variety of cell types with similar representation between rash types (**Fig 5A, Fig S5A-C**). When comparing all clusters in the dataset, we observed a significantly greater proportion of proliferating T cell sin ICI-LP compared to LP (**Fig 2A,B**). However, when we subsetted on T cell clusters to perform a more in-depth comparison of transcriptional states, we noted surprisingly few transcriptional differences in the T cell populations between ICI-LP and LP, likely reflecting the similarity of these two eruptions. DE analysis at the pseudobulk level did show that ICI-LP lesional T cells had higher expression of *AIF1* than LP lesional T cells (**Fig 5B**). Although typically regarded as a macrophage marker, where its expression promotes chemokine and IL-6 production^79^, *AIF1* expression in T cells has been previously reported, and has been shown to increase chemotaxis^80^ and potentiate cytokine secretion^81^. Pathway analysis of T cell clusters showed that ICI-LP T cells exhibit broadly increased JAK-STAT response with an increase in androgen response in Tregs (**Fig 5C-D**).Interestingly, androgen receptor has been reported to enforce *FOXP3* expression, the lineage-defining transcription factor in Tregs^82^. Collectively, these data support a model where there are not dramatic differences in the transcriptional state between the T cell subsets isolated from inflamed skin in ICI-LP vs. LP. Rather, our findings point to more significant differences in fibroblasts and myeloid populations between conditions than T cells.

**Figure 5:**
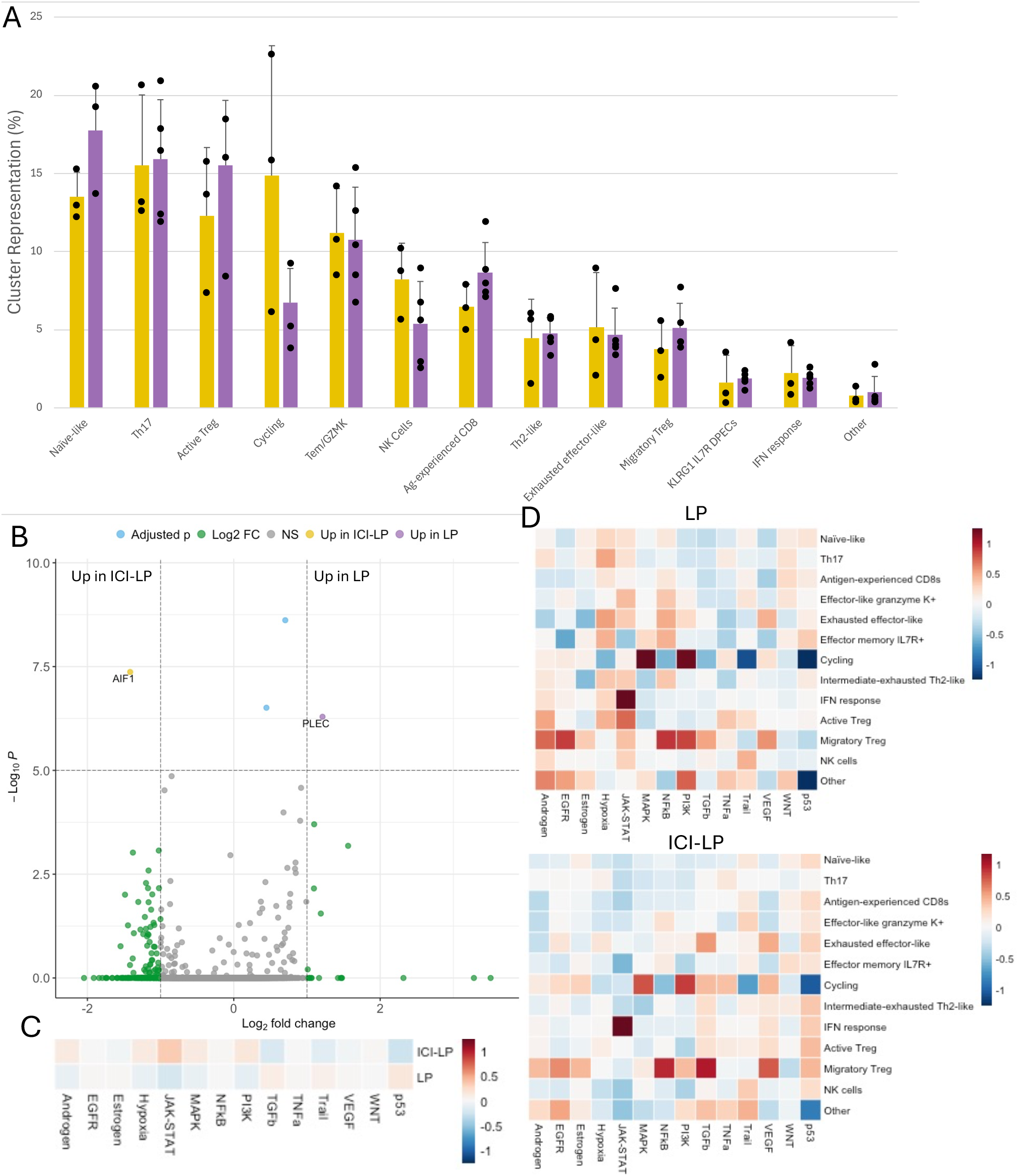
T cell transcriptional phenotype is similar between ICI-LP and LP. (A) UMAP plot of T cell subclustering. (B) Distribution of CD4 and CD8A expression on UMAP. (C) Dot plot of marker genes used in cell type annotation in A. (D) Contribution of UMAP clusters to total composition of T cells, comparing ICI-LP to LP. (E) DiDerentially expressed genes between ICI-LP and LP T cells based on pseudobulking. Threshold for annotation is Log_2_ fold change > 1 or < -1 and -log_10_ p > 5. (F) Comparison of pathway activation in ICI-LP and LP T cells using PROGENy. (G) PROGENy pathway activation by subcluster.

### Spatial profiling reveals localized immune zones in ICI-LP vs LP that show differential inflammatory signatures

Because the single-cell data showed limited divergence in T cell states despite prior reports of mixed Th1-, Th2-, and Th17-associated features in ICI-LP^28,52^, we next asked whether these inflammatory programs might be spatially segregated within lesional skin. We therefore evaluated IL13, IL17A, and IFNG expression in ICI-LP and LP using RNA in situ hybridization (RISH). With this approach, we found expression of all three cytokines within lesional skin. *IFNG* expression was higher than both *IL13* and *IL17A* in LP, whereas *IFNG* exceeded *IL17A* but not *IL13* in ICI-LP samples (**Fig 6A**), in concordance with Mazin *et al*^28^. We considered the possibility that a nonspecific proinflammatory stimulus such as PD-1 blockade may be simultaneously driving multiple arms of immune activation within the same tissue, and that non-spatially-resolved single-cell or pseudobulked analyses may miss smaller niches within lesional skin that would better represent individual local tissue influences on phenotypes of T cells. To assess this possibility, we used spatial transcriptomic analysis of ICI-LP and LP patient samples along with previously-established gene signatures of cytokine responses^77,78,83^. In both ICI-LP (**Fig 6B**) and LP (**Fig 6C**) samples, we indeed found evidence of response to both IL-13 and IL-17A, localized within separate regions, with IL17A response being most evident in the epidermis and IL-13 response focused within the dermis, while IFNγ response was highest at the dermal-epidermal junction. Previous studies have shown AR signaling to promote Treg suppressive activity^84^ while tempering Th17 but not Th2 inflammation^85^, while canonical IGF1 signaling promotes Th17 differentiation^65^. Despite this, evaluation of the localization of IGF1 response within these samples showed a less focused pattern of response, localized primarily within the reticular dermis (**Fig 6B,C**). We sought to confirm this with RISH, and indeed found that all samples evaluated contained IGF1 expression localized primarily within the dermis (**Fig 6D**).

**Figure 6:**
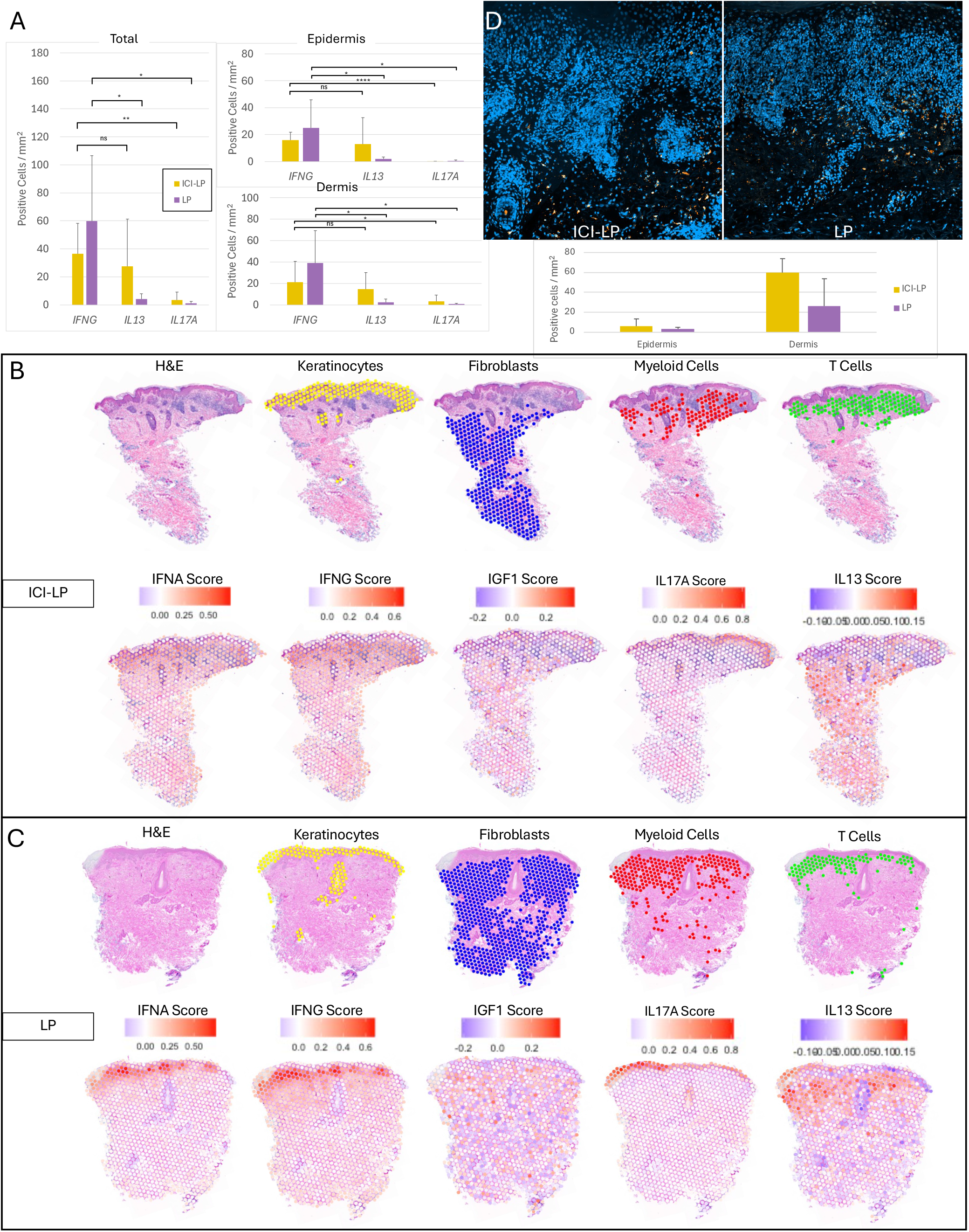
Localized immune zones exist within ICI-LP and LP lesions. (A) RISH analysis of *IFNG, IL13*, and *IL17A* within lesional ICI-LP and LP skin. Left – total section, top right – epidermis, bottom right – dermis. (B) Spatial transcriptomic analysis of LP lesion. Top, cell type identification based on cluster scores derived from scRNAseq analysis. Bottom, cytokine response scores based on signatures established based on *in vitro* cytokine treatment.(C)Spatial transcriptomic analysis of ICI-LP lesion. Top, cell type identification based on cluster scores derived from scRNAseq analysis. Bottom, cytokine response scores based on signatures established based on *in vitro* cytokine treatment. (D) RISH for *IGF1* in ICI-LP and LP lesions. Blue, DAPI; Orange, *IGF1*. Spatial data are representative of at least 5 independent experiments per condition. *, p<0.05; **, p<0.01; ns, not significant.

**Figure 7:**
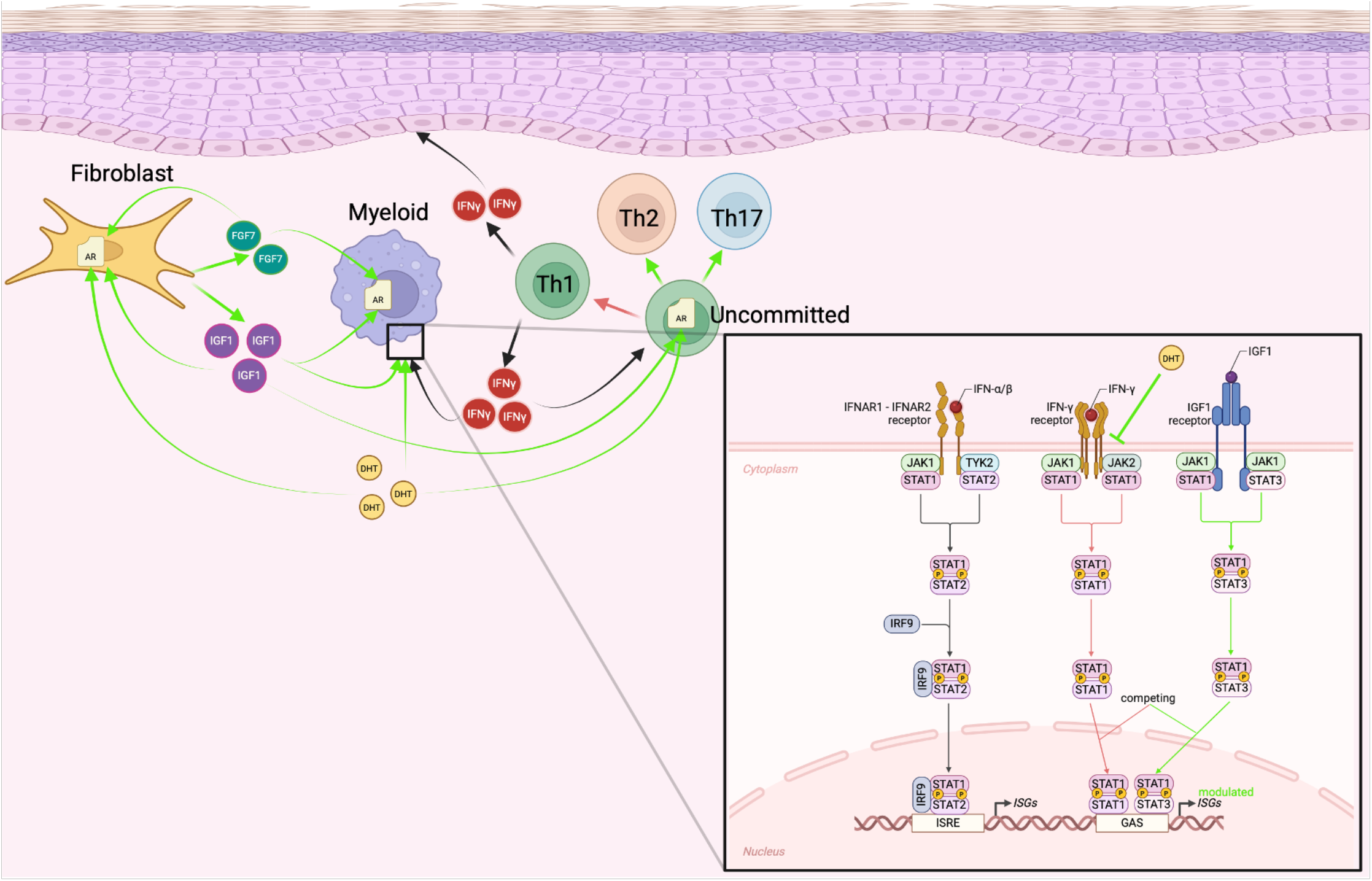
Hypothesized model of IGF1/AR modulation of inflammation in ICI-LP. The dominanr inflammatory process, of Th1/IFN-γ*-mediate*d kertinocyte killing,is modulate in the presence of androgen (here represented by DHT)and fibroblast-secreted IGF.TheseThese mesiatore contribute to a shift in JAK/STAK singnaling, from primarily JAK1/2-driven Stat1 dimers toward JAK1-g\driven STAT1/3 homodimers,leading to modulation of ISG expression profile,ultimately driving diversifiedT-helper fate determination toward Th2 and Th17 phenotypes.This process occurrs with in regional niches within the skin,leading to pocketsof inflamation of disparate phenotypes.

Together, our analyses show that ICI-LP and LP share a broadly similar lichenoid inflammatory framework but diverge most clearly in stromal fibroblast and myeloid compartments. Fibroblasts in ICI-LP were enriched for IGF1/FGF7 and androgen response associated programs, whereas myeloid cells displayed amplified interferon-responsive and JAK-STAT signaling states. By contrast, T cell and keratinocyte compartments were comparatively conserved, and spatial analyses suggested that regional immune compartmentalization contributes to the mixed inflammatory features observed in both conditions.

## Discussion

ICI-induced lichen planus is a particular clinical challenge in oncodermatology because, unlike some other common cutaneous immune-related adverse events, it lacks a well-defined mechanistic framework that could support targeted, steroid-sparing treatment while preserving antitumor immunity. In this study, we compared ICI-LP with spontaneous LP using single-cell and spatial approaches and found that, although the two conditions share a broadly similar lichenoid inflammatory architecture, they diverge most clearly in stromal fibroblast and myeloid compartments rather than in T cells or keratinocytes. Specifically, ICI-LP was characterized by fibroblast enrichment of IGF1, FGF7, and androgen-response-associated programs, together with amplified interferon-responsive and JAK-STAT-active myeloid states. Spatial analyses further suggested that anatomically segregated cytokine-response niches contribute to the mixed inflammatory features observed in both ICI-LP and LP. Tison and colleagues found increased B cells and myeloid cells in ICI-LP alongside relatively fewer CD4^+^ T cells when comparing ICI-LP and LP^86^. Others found ICI-LP to be more likely to present with widespread distribution, with fewer lymphocytes overall but greater exocytosis histologically. Using a 730-gene panel, they found increased evidence of IL12 signaling, overexpression of phagosome genes, and a nonsignificant increase in macrophage representation in ICI-LP versus LP^87^. Mazin et al found a higher CD4/CD8 ratio in ICI-LP than in lichenoid dermatitis, with a ratio of Th2 (GATA3^+^) to Th1 (Tbet^+^) cells more similar to eczematous eruptions than to lichenoid dermatitis^28^, while a separate study found elevated *IL13* and *IL17A* in addition to elevated *IFNG* expression in ICI-LP patient skin when compared to normal controls^52^. Our findings build on these observations by suggesting that the differences between ICI-LP and LP are organized less by wholesale repatterning of lymphocyte states than by selective remodeling of stromal and innate immune compartments. More broadly, this study defines a comparative cellular and spatial framework for distinguishing checkpoint blockade-associated lichenoid inflammation from spontaneous disease, while identifying fibroblast-associated IGF1/androgen signaling as one component of this divergence.

The altered sex distribution of ICI-LP relative to LP motivated our examination of hormone-responsive programs, but our data ultimately point more broadly to tissue microenvironmental differences between these conditions. Within that framework, one of the most striking findings was the identification of IGF1 and FGF7 as among the most elevated transcripts in ICI-LP, with expression localized primarily to fibroblasts and accompanied by enrichment of androgen-response-associated programs. The interplay between androgen and IGF1 signaling is well-recognized^69^. In prostate cancer, both IGF1 and FGF7 have been shown to activate AR^88^, and conversely, AR signaling has been shown to promote IGF1 and FGF7 expression in fibroblasts^89^.Here, too, the expression of IGF1 and FGF7 was primarily in fibroblasts, correlated with evidence of elevated androgen signaling (**Fig 3**). The role of fibroblasts in cutaneous immune responses is an area of rapidly evolving research; prior studies have shown that fibroblasts can maintain a Th2-supporting niche^61^ within the skin, and a recent fibroblast atlas delineated features of “disease-adapted” fibroblasts that were found in lichenoid skin disease^62^ and shared features with populations we identified (**Fig 3E**). These observations support the idea that fibroblasts are not simply structural bystanders in lichenoid inflammation but may help shape the inflammatory context in which checkpoint blockade-associated disease develops.

The immunologic consequences of androgen- and IGF1-associated signaling in skin are likely complex and context dependent. AR signaling has been shown to be broadly suppressive of Th1-supported cytotoxic responses^36-38^, while IGF1 signaling has been shown to promote T cell-mediated inflammation in a variety of contexts^90-93^, including in oral lichen planus^91^. However, IGF1 exposure also desensitizes T cells to IFN-γ *through* internalization of IFN-γR2, leading to a net decrease in STAT1 activation^94^. In its place, STAT3 and STAT5 activation supports the development of a Th17 phenotype^65^. Notably, we did see a Th17 population in both ICI-LP and LP (**Fig 5D**), although this was similar in both conditions. In myeloid cells, IGF1 is also a pro-inflammatory influence, increasing production of IL6, IL1B, and TNFa while supporting macrophage activation^90^. IGF1 signaling can also promote M2 polarization of macrophages, which support a Th2 inflammatory phenotype^95,96^. Loss of signaling through the IGF1 receptor, on the other hand, has been implicated in the loss of peripheral tolerance in autoimmune disease^97^. Taken together with our data, this literature suggests that IGF1-associated fibroblast programs may influence the balance of inflammatory signals within lesional skin and thereby contribute to the heterogeneous immune phenotypes reported in both LP and ICI-LP. Our spatial analyses are consistent with this possibility, as they revealed anatomically segregated IFNγ-, IL-17A-, and IL-13-response niches within individual lesions rather than a single uniform inflammatory state. This regional organization may help explain why prior studies, and even different analytic modalities within the same study, identify overlapping but not identical Th1-, Th2-, and Th17-associated features.

A second important finding was that myeloid populations, more than T cells, carried the strongest differential interferon-responsive and JAK-STAT-active programs in ICI-LP. This is notable because lichenoid inflammation is often conceptualized primarily through T cell-mediated keratinocyte injury. In our data, however, T cell states were comparatively similar between ICI-LP and LP, whereas myeloid cells showed broad enrichment of interferon-associated programs across multiple subpopulations. These findings suggest that the tissue context surrounding T cell-mediated injury differs substantially between checkpoint blockade-associated and spontaneous lichenoid disease, and that innate immune amplification may be a major feature of ICI-LP pathogenesis.

The current study is limited by its sample size, the restriction to male patients, and the lack of a manipulable model system, and additional work will be necessary to identify the generalizability and clinical significance of these findings. Our findings outline a potential role of androgen signaling to the immunophenotype of ICI-LP as compared to LP. This may contribute to the observed disparity in incidence by sex between these conditions if an AR signaling deficit is a necessary precondition for the development of spontaneous LP but not necessary during ICI therapy, although this is unlikely to be a complete explanation. Our study does not examine the role of other sex hormones, or of other noted mediators of sex-based differences in immunity such as XIST^98^ or VGLL3^99^. While additional study will be needed to disentangle these factors, as well as the specifics of IGF1’s role in the immunophenotypes of ICI-LP and LP, the fact that pharmacologic interventions already exist to modulate IGF1^100^, androgen^101^, IL4/13^102^, and IL17^103^ signaling underscores the potential for rapid translation once these features are clearly understood.

## Materials and Methods

### Multiplexed Fluorescent RNA in situ Hybridization

Five-micron paraffin-embedded sections were stained with ACD’s RNAscope Multiplex Fluorescent Detection reagents V2 reagents (ACD #323110) according to manufacturer’s instructions. All slides underwent deparaffinization, pretreatment to unmask target RNA and permeabilize cells, probe hybridization, and signal amplification with Akoya opal fluorophores. The probe sets and their corresponding fluorophores were Opal 520 1:1500 dilution (# FP1487001KT), Opal 570 1:1000 dilution (#FP1488001KT), and Opal 690 1:1000 dilution (# FP1497001KT). The sections were then counter stained with DAPI and mounted with ProLong Gold Antifade reagent (Invitrogen #P36930). Positive and Negative controls were generated using ACD’s RNAscope positive and negative probesets. Slides were imaged on a Zeiss Axioscan 7 Microscope Slide Scanner at 20X. Scanning was performed using ZEN Blue 3.7 software using a Colibri7 LED light source, Plan-Apo10 × 0.45 NA objective, and an Orca Flash4.0 C13440 camera. Images were pre-processed in ZEN Blue 3.9 (Zeiss). Images with stitching artifacts were re-stitched using the ‘fuse tiles’ function. Images were analyzed in Arivis Pro 4.4 (Zeiss) using a custom pipeline as follows. First, nuclei were segmented based on the DAPI channel using a custom Cellpose2.0 model, trained via the Cellpose Python GUI. From these nuclei, cell boundaries were estimated using region growing with a max dilation of 2.5 µm. Then, fluorescent puncta were detected using the ‘Blob Finder’ algorithm. Diameter (2um), split sensitivity (67%), and normalization ranges (0-16383) were applied consistently across all images. Blob Finder probability thresholds were set based on the signal-to-noise ratio of each mRNA probe. Autofluorescent artifacts, present across all channels in human FFPE skin, were filtered out on a per-image basis using multi-channel intensity thresholds to account for inter-patient autofluorescence variability. Within the estimated cell boundaries, the ratio of fluorescent puncta area to cell area was calculated, and cells with a ratio >0.035 were considered positive for labeled mRNA. Cell counts were normalized to 1 mm^2^ based on the analyzed area.

### Immunohistochemistry

Sample Processing: Androgen Receptor (Cell Marque #200R-17) staining was completed by OHSU’s Histopathology Shared Resource with prostate tissue utilized as the positive control.

IGF1-R (R&D Systems # MAB391) and ISG15 (Abcam # ab285367) staining were done with MCF7 cells as a positive control. For the coverslips, we grew MCF7 cells in 75cm flasks and then transferred ∼200,000 cells into each well of a 6 well plate that contained coverslips. Once cells were adequately spread onto the coverslip (∼24-48 hours), they were pre-fixed by adding 500 µL of Thermo Scientific 4% Paraformaldehyde Fixative Solution (#J61899.AP) directly into the culture medium. After 2 minutes, we replaced the pre-fixation culture medium with 300-400 µL of 2% Formaldehyde Fixative Solution and incubate, for 20 minutes at room temperature. Then the coverslips were rinsed 3X with 1X-PBS and stored at 4c.

We used the Thermo Scientific Gemini AS slide stainer to deparaffinize the FFPE slides, then we performed Heat Induced Antigen Retrieval (HIER) on the tissue sections mounted on charged glass microscope slides at 97c for 20 minutes with Epredia HIER Buffer H pH 9.0 (#TA-135-HBH). We did a 1-hour incubation for both the IGF1-R and ISG15 antibodies, which was followed by the Bio SB PolyDetector Plus DAB HRP Brown Immunohistochemistry (IHC) detection system (#BSB 0263). We counterstained with Epredia Hematoxylin 1 (#88018) for 10 seconds then rinsed the slides with 1X PBS and used Epredia Aqua-Mount Media (#14-390-5).

### Spatial Transcriptomics

Sample Processing: Obtained 5-miron sections from either MD Anderson or OHSU Dermatopathology. Tissue sections were deparaffinized, then stained with hematoxylin (3 min), bluing reagent (1 min), then eosin (45 seconds). Brightfield images were generated on the Zeiss Axioscan 7 Microscope slide scanner at 20X. After imaging, the coverslip was removed as per 10X Visium CytAssist Spatial Gene Expression for FFPE – Deparaffinization, H&E Staining, Imaging & Decrosslinking demonstrated protocol (CG000520) and the slides were inserted into the Visium CytAssist moveable cassette frame and gasket (#3000813, #300814 & 300816) to ensure there were no leaks when adding the reagents. The slides were then destained and decrosslinked to release RNA that was isolated by the formalin fixation. The 10X Visium Cytassist Spatial Gene Expression user guide (CG000495) was followed starting with probe hybridization overnight for 17 hours. Post hybridization wash, probe ligation, and post ligation wash followed to bridge the junction between the probe pairs and the RNA. The probe release and capture are then conducted in the Visium CytAssist instrument where the ligation products are released from the tissue and captured on the visium slide. The ligation products are then extended by the addition of the UMI, spatial barcode, and partial read 1 primer. These products are amplified then cleaned up using SPRIselect. This pre-amplification material is utilized for qPCR to determine the ideal cycle number for the sample index PCR. We added +2 cycles to the CQ values determined from the qPCR as per 10X user guides’ instructions to use for the SI-index PCR’s cycle number. The resulting library was cleaned up by SPRIselect and then assessed on a bioanalyzer using the DNA high sensitivity assay and sequenced on an Illumina NovaSeq 6000 by OHSU’s Integrated Genomics Lab.

Sequence Alignment and Annotation: The cytassist and histology image files were utilized to align the fiducial frames on Loupe Browser 8 (10X Genomics) then exported as a JSON for SpaceRanger. Sequencing output was processed using SpaceRanger software (10X Genomics) by OHSU’s Integrated Genomics Lab. Then Illumina’s bcl-convert (before Jan 2025, we use bcl2fastq) was used to convert basecalling data to fastq and to demultiplex. The Space Ranger count function was used to align reads and calculate counts based on the human reference genome (GRCh38) and then align microscopic slide images and transcriptomes to generate barcode/UMI counts and feature spot matrices.

### Single Cell RNA-sequencing

Sample Processing: We followed 10X Genomics demonstrated protocol CG000632 to isolate nuclei from the FFPE tissue sections. Each sample started with 6 scrolls, which we then dissociated following the pestle protocol. We had three tubes for each sample with two scrolls in each tube, which were centrifuged after all three of the ten-minute xylene washes for 5 min at 850 rcf. The three sample tubes were then combined during the addition of the PBS step. After the 45-minute dissociation incubation, the samples were aspirated by a 23G x 5/8” hypodermic needle to improve cell recovery. We then followed 10X Chromium Fixed RNA Profiling Reagent Kits – Multiplexed user guide (CG000527) to perform probe hybridization where the left-hand side and right-hand side probe pairs would hybridize to their complimentary RNA in an overnight incubation (17.5 hours). We then pooled 4 different samples together; each sample would get a barcode (BC001-BC004) with a targeted cell recovery of 40,000 (10,000 per sample). GEMs were generated by combining the barcoded gel beads, the master mix containing the cell suspension and post-hybridization resuspension buffer, and partitioning oil into the Chromium Next GEM Chip Q which was subsequently run on the Chromium iX machine. The GEMs mixture was then incubated, and a recovery step was performed. A PCR is done to pre-amplify the ligated products, which are cleaned up by SPRIselect. This pre-amplificated product underwent sample index PCR which generated the final library. Finally, the product is run on the 2100 Bioanalyzer instrument using the DNA high sensitivity assay and sequenced on an Illumina NovaSeq 6000 by OHSU’s Integrated Genomics Lab.

### Transcriptomics Data Analysis

Preprocessed gene expression matrices were loaded into Seurat^104^ (version 5.1.0) from hdf5 files using Seurat’s Read10X_h5() function. Raw hdf5 files, filtered hdf5 files, and clusters CSV files were used as input to SoupX^105^ (version 1.6.2) to remove reads corresponding to ambient (cell-free) RNA using default parameters. Cells with more than 15% mitochondrial genes and less than 200 unique genes detected were removed from each dataset. Normalization was performed with SCTransform and samples were integrated using CCA dimensionality reduction. Dimensionality reduction was performed on the integrated Seurat object (PCA, UMAP using the top 20 dimensions), and unsupervised clustering was performed using FindNeighbors() function with the top 20 dimensions, and FindClusters() function. Optimal resolution was set at 0.15, and clusters were annotated using canonical marker genes and marker genes from the literature.

Pseudobulking for differential expression (DE) analysis was performed using Seruat’s AggregateExpression() function, with DE testing using the DESeq2 test. Cytokine response scores were calculated using Seurat’s AddModuleScore() function. Topic modeling was performed using the TITAN package^63^, with a number of topics chosen based on perplexity using TITAN’s LDAelbowplot() function. Pathway analysis was performed using PROGENy^59^ via decoupleR^106^.

## Acknowledgements

The authors gratefully acknowledge the support of the Melanoma Research Foundation, the National Center for the Advancement of Translational Science, the Collins Medical Trust, and the Dermatology Foundation. The research reported in this publication used computational infrastructure supported by the Office of Research Infrastructure Programs, Office of the Director, of the National Institutes of Health under Award Number S10OD034224. The content is solely the responsibility of the authors and does not necessarily represent the official views of the National Institutes of Health.

