## Supplementary figures and images for "IGF1 modulates lesional skin inflammation in checkpoint inhibitor-induced lichen planus"

### Supplemental Materials

**Supplemental**

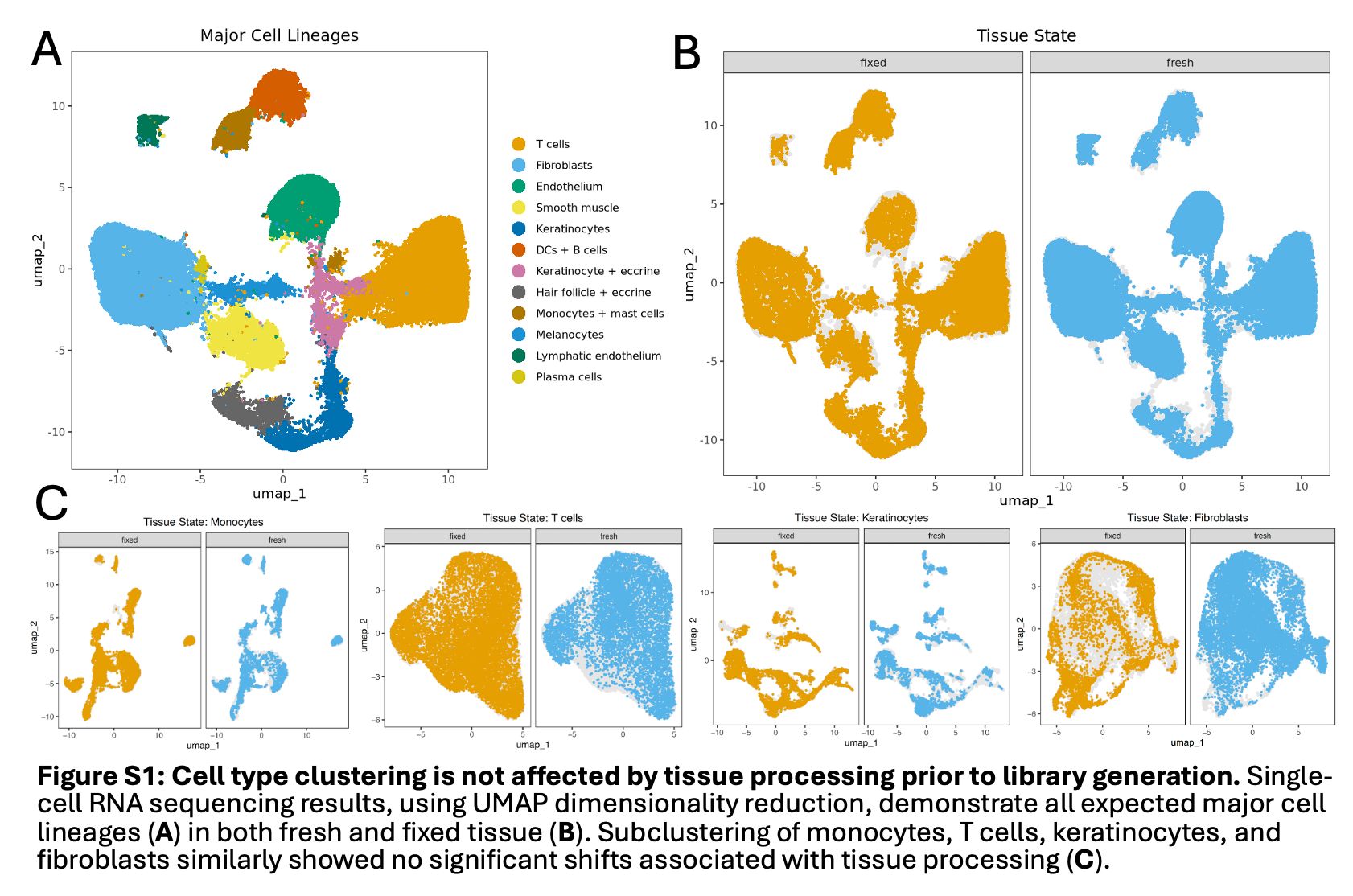

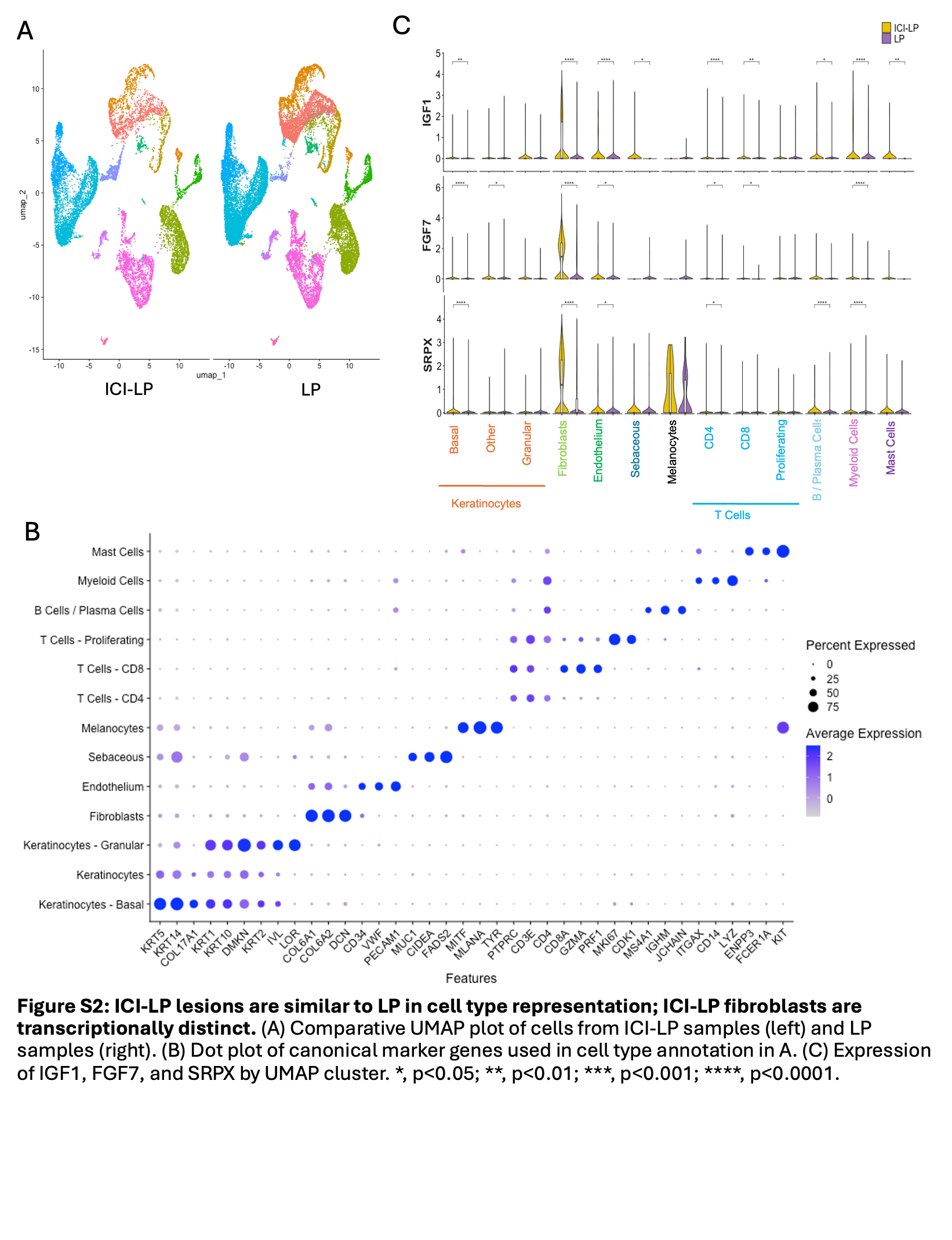

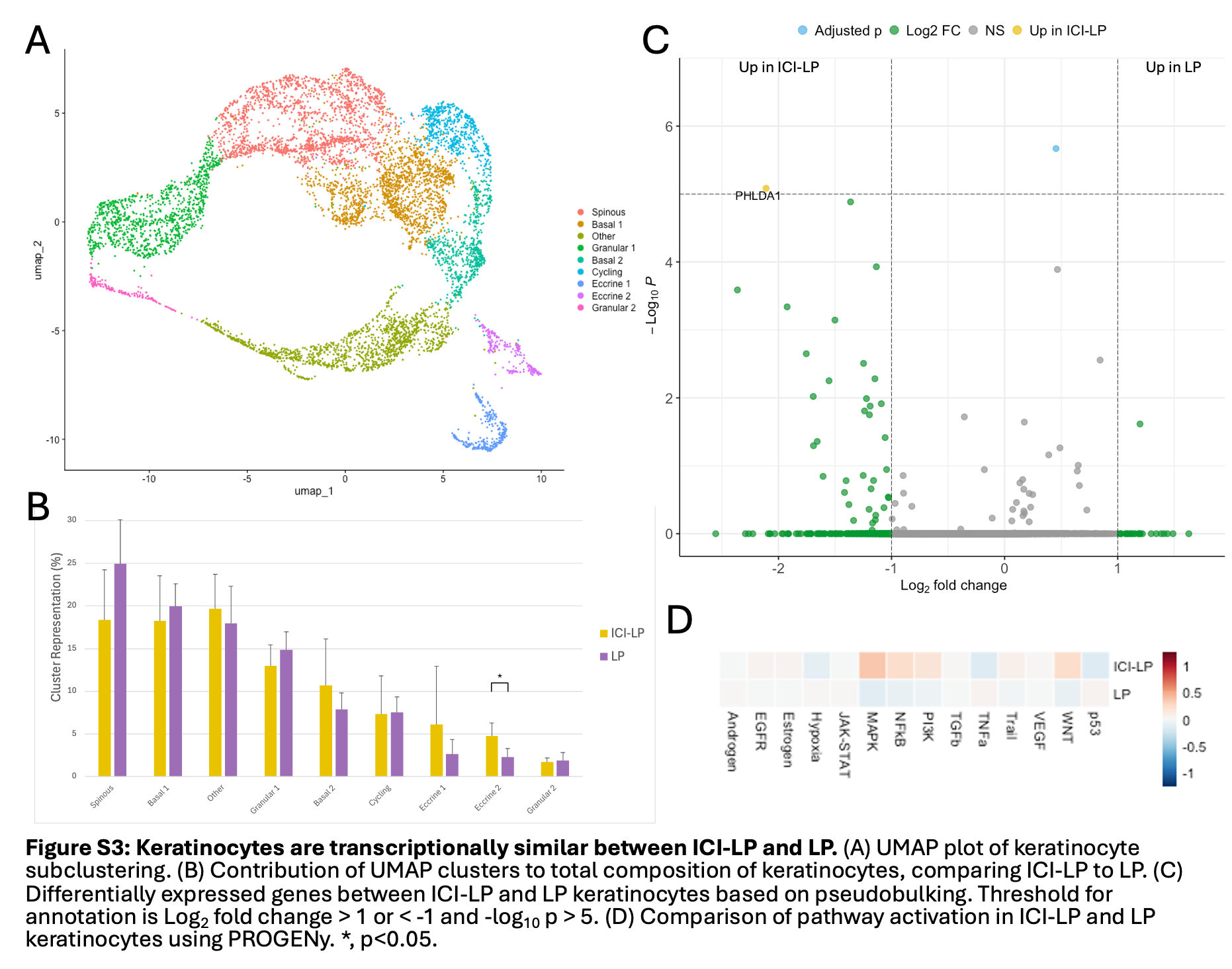

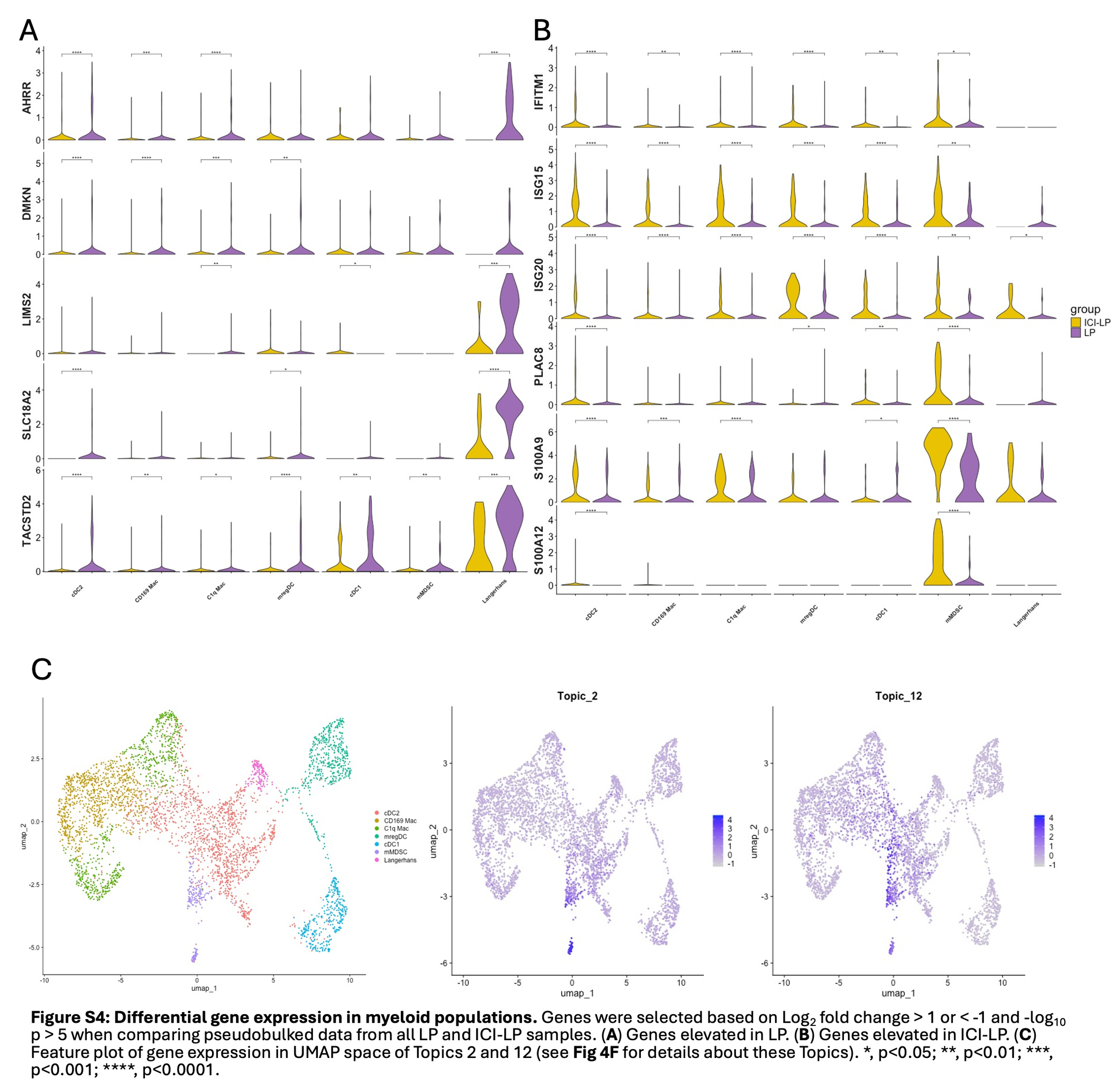

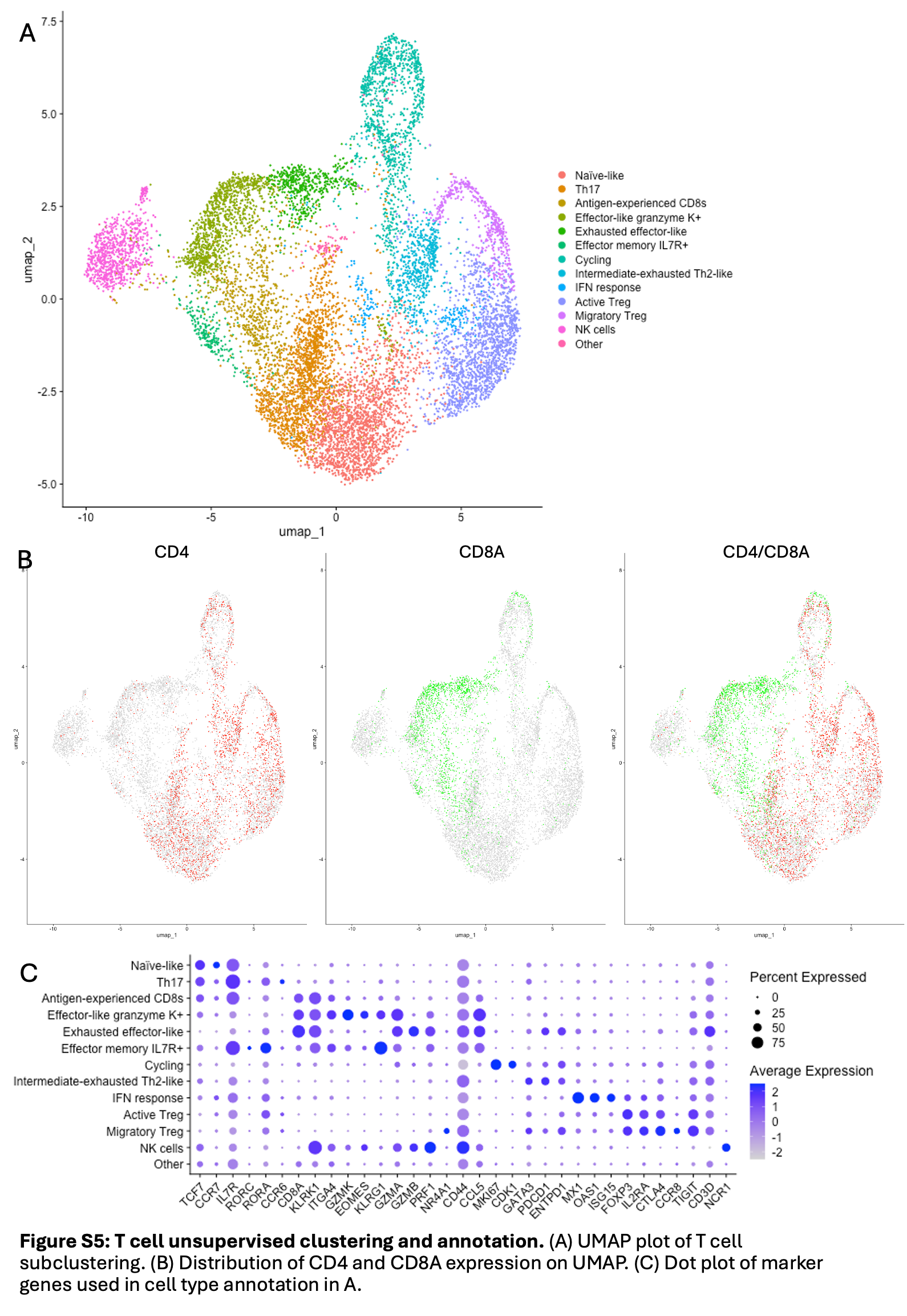

**Table S1: Patient Characteristics**
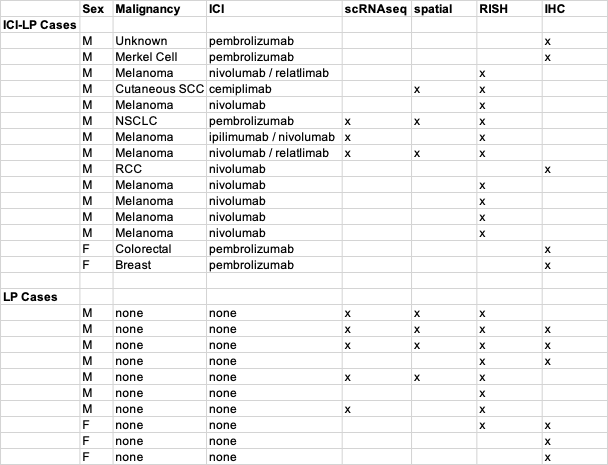
